# Binocular disparity can augment the capacity of vision without affecting subjective experience of depth

**DOI:** 10.1101/222034

**Authors:** Henry Railo, Joni Saastamoinen, Sipi Kylmälä, Aapo Peltola

## Abstract

Binocular disparity results in a tangible subjective experience of three-dimensional world, but whether disparity also augments objective perceptual performance remains debated. We hypothesized that the improved coding of depth enabled by binocular disparity allows participants to individuate more object at a glance as the objects can be more efficiently differentiated from each other and the background. We asked participants to enumerate objects in briefly presented naturalistic (Experiment 1) and artificial (Experiment 2) scenes in immersive virtual reality. This type of enumeration task yields well-documented capacity limits where up to 3-4 items can be enumerated rapidly and accurately, known as subitizing. Our results show that although binocular disparity did not yield a large general improvement in enumeration accuracy or reaction times, it improved participants’ ability to process the items right after the limit of perceptual capacity. Binocular disparity also sped-up response times by 27 ms on average when artificial stimuli (cubes) were used. Interestingly, the influence of disparity on subjectively experienced depth revealed a clearly different pattern than the influence of disparity on objective performance. This suggests that the functional and subjective sides of stereopsis can be dissociated. Altogether our results suggest that binocular disparity may increase the number of items the visual system can simultaneously process. This may help animals to better resolve and track objects in complex, cluttered visual environments.

---

Humans and many other animals have two forward facing eyes and a large overlap between their visual fields 1. One of the major benefits of this is stereopsis: integration of the slightly different viewpoints of the two eyes (binocular disparity) results in tangible subjective sensation of three-dimensional visual space, and accurate coding of depth 2,3,4. Binocular disparity improves eye-hand coordination 5,6,7, and visual recognition of singular items when simple stimulus displays are used 8,9,10,11. Whether disparity also improves the segmentation of objects from background in naturalistic settings where multiple objects are present remains open 12. Specifically, we were interested in examining if binocular disparity improves the recognition of multiple objects in naturalistic settings. We hypothesized that because disparity allows differentiating between different objects in depth it enables faster and more accurate individuation of the objects. This could enable more efficient visually-guided behavior, and hence improve the individual’s chances of survival.

Stereopsis is one of the defining features of primates 13, and it has been linked to the expansion of brain structures that process visual information 14. Early theories explained the emergence of frontal eyes as adaptations to arboreal locomotion because movement in arboreal environment (for instance, when leaping between tree branches) requires accurate judgement of distances 15,16,17. The arboreal theory of stereopsis has been challenged as, for instance, many non-primate species (such as the squirrel) are capable of skilled arboreal locomotion despite the lack of a binocular visual field 18 suggesting that arboreal locomotion alone does not explain the evolution of frontal eyes. According to the alternative visual predation hypothesis, stereopsis is an adaptation that enables animals to better visually locate and seize prey without moving (i.e. without motion parallax; 18,19). Similarly, the camouflage-breaking hypothesis of stereopsis argues that stereopsis enables animals to detect camouflaged prey 20. The visual predation theory was later elaborated to concern especially nocturnal animals as a large binocular visual field meant that the animals could better detect objects in dark environments 21. However, the visual predation theory is undermined by the fact that some excellent predators do not have frontal eyes, and early nocturnal primates may have detected their prey (mainly crawling insects) using olfaction or hearing 22,23,24. Moreover, these smaller primates were largely omnivorous, eating lots of plant material in addition to insects, which lead Sussman (22, p. 219) to argue “that the explanation [for orbital convergence] will be found in adaptations providing the fine discrimination needed to exploit the small food items available on the newly diversifying flowering plants”. Establishing the possible functional advantage provided by visual disparity is crucial to understand what factors may have driven the evolution of frontal eyes.

Random dot stereograms illustrate how stereopsis may enhance perception and enable the detection of otherwise hidden objects 20, but they represent extremely simplified cases: all other depth cues (than binocular disparity) have been removed, and the object is placed in front of a flat background surface. In real life, the objects to be detected are embedded in scenes that include various depth cues (shadows, occlusion, object size etc.), and the scenes are often cluttered with other objects that occupy different depths. To understand if binocular disparity improves the speed and accuracy of object processing, it is important to use stimuli that more closely resemble natural environments. There is anecdotal evidence that stereopsis helps observers to detect objects in realistic scenes as aerial stereographic photographs (with greatly exaggerated visual disparity) have been used to detect camouflaged objects since the first world war 25. Caziot and Backus reported that binocular disparity improved the recognition of stimuli displayed in front of a flat textured background 9. Because this facilitation was observed even when the disparity did not coincide with the object’s luminance contour, the authors concluded that binocular disparity facilitated allocation of attention towards the object. McKee et al. 26 has shown that visual disparity aids the detection of a moving target dot that is flanked by randomly moving distractor dots that occupy a different plane in depth, but crucially when the moving target is presented in the midst of randomly moving distractors, disparity provides negligible help 26. Valsecchi et al. 12 found that binocular disparity did not improve the accuracy of recognizing artificial or naturalistic scenes when monocular cues were available to provide information about the structure of the scene. However, McKee and Taylor 27 showed that when real stimuli (instead of computer generated stimuli) were used, binocular depth discrimination thresholds of two adjacent objects were clearly superior to monocular thresholds.

To study if binocular disparity improves figure-ground segmentation in naturalistic environments, we asked participants to rapidly detect and enumerate individuals presented in an immersive virtual environment. Enumeration reveals well-documented limits in visual capacity: when 1-3 items are presented, enumeration is fast, accurate and relatively independent of number – a process termed subitizing. Larger sets decrease enumeration accuracy and increase response times 28,29. The number of items that can be “subitized” is considered to mark the capacity of items that can be individuated and differentiated from the background and each other in a single glance 30,28,31,32. If binocular disparity increases the capacity of perception, enumeration should become faster and more accurate when compared to the condition where visual disparity has been removed (the same image is presented to both eyes). Figure 1 presents an overview of the present experiment and the hypotheses. One can hypothesize that the information added by binocular disparity enhances performance across different set sizes (Fig. 1B, left), for example, because disparity enables more efficient figure-ground segregation because the items can be resolved in depth when disparity is available. The influence of disparity could be limited to smaller set sizes, i.e. those items that the participants can efficiently process (Fig. 1B, middle). Third, it could be assumed that binocular disparity allows participants to augment the capacity of perception by enabling them to subitize larger set sizes (Fig. 1B, right). Although Fig. 1B only presents hypothesis concerning enumeration accuracy, similar hypotheses can be made also about reaction times.

**Figure 1:**
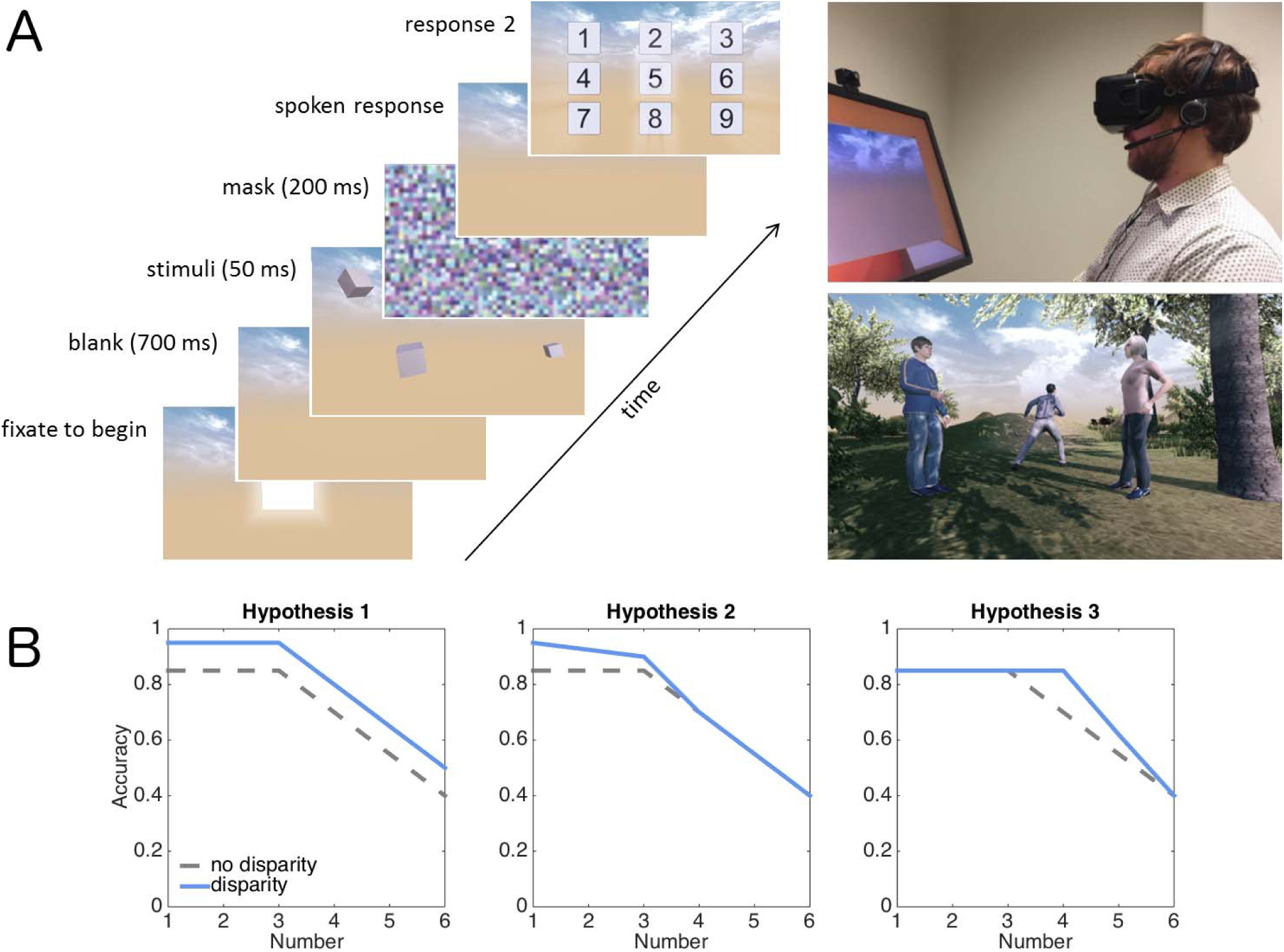
A) Task and stimuli. Schematic presentation of a single trial with cube stimuli (left; Experiment 2), the experimental set up (top right), and an example of a scene stimulus (Experiment 1). In the experiments, the stimuli (1-6 objects) were always presented binocularly and disparity was manipulated. In one condition the participants were asked to enumerate the individuals in the scene and in the other condition the participants rated the strength of their experience of depth. B) Hypotheses concerning enumeration accuracy.

**Figure 2:**
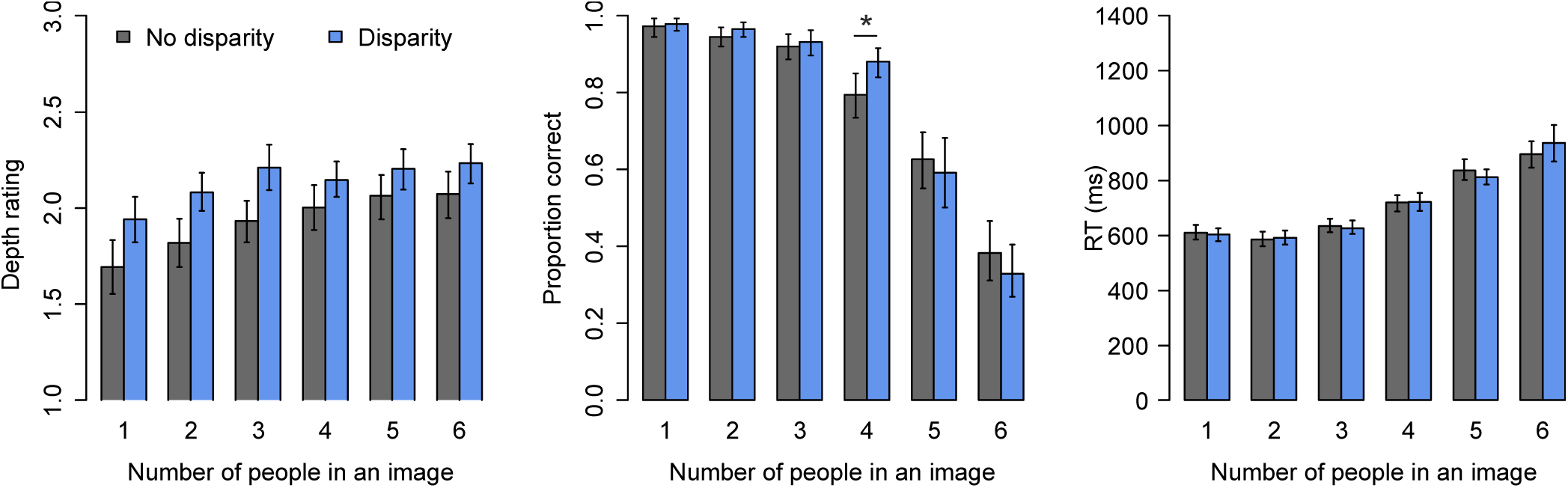
Results of Experiment 1. Error bars show 95% CIs.

A further interesting open question concerns to what degree the possible improvement in objective visual functioning (e.g. lower detection thresholds) due to binocular disparity is mirrored by changes in subjective, consciously experienced perception of depth. While many classical studies use the term stereopsis to refer both to the subjective (qualitative) experience of immersive depth and the objective (quantitative) coding of depth, Vishwanath 3 has argued that the two senses of the term stereopsis should be distinguished. Consequently, the effects of binocular disparity can be measured in two ways. First, one can examine how binocular disparity influences subjective experience of depth, for instance, by asking participants to rate the strength of their subjective experience of depth (such rating scales are commonly used in studies of conscious vision, see e.g. 33,34). Second, one can measure how binocular disparity affects objective, visually-guided behavior (e.g. recognition accuracy and reaction times). Studies suggest that these two senses of the term stereopsis can be dissociated: Subjective sensation of depth can emerge without binocular disparity 35, and disparity can influence behavior in the absence of subjective sensation of depth 36,37. Concerning the present study, we asked whether binocular disparity has similar effects on the subjective and objective sides of stereopsis. Or can binocular disparity increase the participants’ subjective experience of depth without affecting their objective visual performance, or vice versa? Typically studies only measure objective visual performance and assume that binocular disparity also enhances the subjectively experienced impression of depth.

The present study employed very brief presentation times (50 msec) combined with backward masking to focus on the contribution of information contained in visual disparity per se. Such brief stimulus durations minimize the contributions from motion parallax (if the person moves his head), but also limits the strength of subjective perception of depth 38. However, if visual disparity contributes to the speed and accuracy of segmenting objects from the background, the effect should be observed even with short presentation times that may be insufficient to produce a strong subjective sensation of depth. The amount of visual disparity was also restricted in the present study to minimize the physical difference between the disparity vs. no-disparity conditions (see Methods). This meant that from the participant’s point-of-view the disparity vs. no-disparity conditions looked very similar (none of the participants reported spontaneously perceiving a change in disparity during the experiment).

## Method

### Participants

Twenty-nine participants took part in Experiment **1** (age range **1**8-34, 23 females), and 31 participants to Experiment 2 (age range **1**9-35, 26 females). All participants reported having normal or corrected-to-normal vision, and reported that they did not have deficits in stereo vision (e.g. strabismus). Participants gave informed consent to participate the study, the study was conducted in accordance with the Declaration of Helsinki, and the ethics committee of the hospital district of South-West Finland approved the study.

### Stimuli and procedure

Participants were presented with stereoscopic and non-stereoscopic stimuli in immersive virtual reality using the Oculus Rift DK2 headset. The two screens of the headset have a resolution of 960 × 1080 per eye (75 Hz refresh rate), and the eyes are focused on the screens with lenses whose focal distance is 1.3 meters. Total field of view is 100°. Immersive virtual reality refers to the fact that the participant feels being present in the virtual world. The world is immersive in the sense that it surrounds the participant and by turning her head the participant can look around in the virtual world. The Oculus Rift was running on a computer (3.50 GHz, 16 GB RAM) with a NVIDIA GeForce GTX 980 graphics card. The camera that tracks the position and orientation of the headset was placed roughly on the table roughly 1 meter from the participant. The distance between the two lenses which focus the viewers gaze of the screens in the headset was adjusted for individual participants to match their IPD.

In Experiment 1 the stimuli were natural scenes that contained 1–6 humans (see Fig. 1). The humans were generated using an open source Make Human software (http://www.makehumancommunity.org). They were distributed around the scene, they never overlapped with each other, and at least their head and torso were clearly visible. All of the humans were roughly at the same height level as the camera, at an average distance of about 3.5 meters (SD = 1.6) from the camera in the virtual reality world. This distance mainly chosen to fit up to six humans in one view. When the scenes contained multiple humans, the objects were scattered around the field of view, although sometimes they also formed sub-groups. The distance of the humans from the virtual camera and the positions of the humans was similar at each number. The scenes, which were generated using various “prefabs” of the Unity software, contained trees, bushes or rocks, but never any artificial, man-made objects. The humans were positioned directly in front of the camera. In Experiment 2 the stimuli were randomly oriented cubes presented at pseudo-random positions in front of the participant on sky background (same as in Experiment 1). The average distance of the cubes relative to the camera was similar as in Experiment 1. The color of the cubes was constant within a scene, but varied across scenes.

The experiment and the stimuli were made using the game engine Unity (5.4.0f3). The images presented to the left and right eye screens in the Oculus Rift were determined by two cameras placed in the virtual world constructed by Unity. We used the default settings: the distance between the left and right virtual cameras was 64 mm, and they were parallel to each other, that is, the cameras were not angled inward like eyes typically are. While this means that while the differences between the viewpoints of the two eyes is reduced, the scenes with and without visual disparity look very similar (the manipulation of disparity was not spontaneously noticed by the participants). This is important as strong variation in disparity could draw the participants’ attention and influence behavior indirectly (note that disparity and no-disparity trials were presented randomly within experimental blocks). Furthermore, stronger than normal disparity could produce nausea in some participants. The inter-camera distance of 64 mm corresponds to the average IPD distance in males. This means that the inter-camera distance is thus higher than the average IPD in females. Subsequent studies should consider adjusting inter-camera distance individually. In the “No disparity” condition the image of the left virtual camera was presented to both left and right eyes of the participant.

As shown in Figure 1, each trial began with the presentation of a light gray square positioned directly in front of the participants. When the participant oriented towards the square for 1 second it turned to a darker shade and triggered the presentation of the scene 700 msec later. Each scene was presented for 50 msec, and it was followed by a flat (i.e. 2D) pixelated mask image for 200 msec. In the enumeration task the participant was asked to say out loud the number of humans presented as quickly as possible. The participants wore Sennheiser PC-8 headphones with a microphone. Reaction times relative the onset of the scene were recorded. After the vocal response numbers 1–9 were presented on a 3 × 3 grid of squares in the virtual reality. The participant selected the number she had spoken by orienting towards it in the virtual reality. The answer was selected when the participant kept orienting towards the number for at least **1** second. The second response was collected like this to induce a more complete sensation of immersion and because keyboard responses without visual feedback might be challenging for some participants. In the subjective experience of depth task the participant was asked to rate the strength of the subjective experience of depth and three-dimensionality on a scale from one to three. The highest alternative denoted a strong sense three-dimensionality, whereas the lowest alternative meant that the participant experienced a relatively flat, 2D world with very little sensation of depth. The middle alternative referred to medium strength experience of depth. We also emphasized to the participant that this task was not related to the apparent quality of the graphics, nor was there a correct answer to the question. Previous studies have shown that participants can assess the strength of experienced depth associated with stereoscopic stimuli independently of other visual attributes (e.g. image quality) 39,40. The participant was told that she did not have to make a speeded response. The answer was given first by speaking to a microphone and then selecting the corresponding response by orienting to it in the virtual reality.

Each participant performed two blocks of 60 trials of the enumeration and two blocks of 60 trials of the depth rating task. The order of the tasks was counterbalanced across participants. Each participant saw each stimulus once, and across participants each stimulus appeared equally often in the enumeration and in the depth rating tasks. Ten practice trials were completed before each task.

### Statistical analysis

Data were analyzed in R statistical software (version 3.**1**.2; 4**1**) to assess how the manipulated factors (disparity and number) influenced response times and enumeration accuracy. Single-trial depth rating and reaction time data was analyzed using linear mixed-effects models 42, and accuracy data with logit mixed-effects models 43 with binomial probability distributions (using the lme4 package; 44). Mixed-effects models enable taking into account individual differences (e.g., in the speed of responding) in the model, in addition to group-level effects. In the mixed-effect models, the number of items (**1**–6), binocular disparity (disparity vs. no disparity), and their interaction were added as fixed-effects factors. In addition, the reaction time of a previous trial was included to reaction time models, and the depth rating of a previous trial was included to the depth rating analyzes. These were included in the model to control for trial-history effects (e.g. 45,46) to obtain more precise estimates of the main factors (disparity and number). Both of these continuous variables were centered. Participant-wise variation was modelled with random-effects including individual intercepts and slopes per participant (for both disparity and number conditions). We used the most complex random-effect structure with which the model still converged. Because enumeration data is known to contain a breakpoint around number 2-4, the data was fitted with two-segment linear models in addition to standard one segment models (for formal description, see 29). We fitted three alternative two-segment models where the length of the first segment ranged numbers **1**–2, **1**–3, or **1**–4. These were compared against the one segment model with a corresponding random-effect structure. Model fits were evaluated using the Akaike Information Criterion, which weighs the goodness of fit by the complexity of the model. Different models were compared using likelihood-ratio tests. The best fitting model was pruned by removing nonsignificant regressors (t < |2|). The pruned models are visualized in the figures. We do not report p values because there is no standardized way to calculate the degrees of freedom and the corresponding p values in mixed-effects models 42. The statistical significance of an effect in regression models can be checked from the confidence intervals (CI) of the regression coefficient: whenever the value zero is included in the CI, the effect is not considered statistically significant. Error bars in all the figures are 95% CIs calculated from **1**000 bootstrap samples.

Datasets and the analysis scripts can be downloaded at the Open Science Framework (https://osf.io/9b3h8/).

## Results

### Experiment 1: Natural scenes

The results of Experiment 1 where the participants viewed natural scenes are presented in Figure At group level, subjective depth ratings decreased .20 units on average when binocular disparity was removed (β = −0.20, CI= [−0.25, −0.15], t = 8.5). Moreover, depth ratings increased with number (β = 0.065, CI = [0.043, 0.087], t = 5.9). The interaction between number and disparity condition was not statistically significant, and was thus left out of the model. Finally, the depth rating of previous trial predicted ratings of depth (β = 0.089, CI = [0.058, 0.12], t = 5.6). Participant-wise random-effect variation related to intercept was SD = 0.26, and related to number was 0.063.

Enumeration accuracy was best fitted with a two segment model (χ^2^ = 12.1, df = 4, p = .016): Accuracy remained high for the first three items (subitizing), after which it declined rapidly. Accuracy was slightly lower without binocular disparity during the first segment (OR = 0.66, CI = [0.49, 0.90], Z = −2.61). Accuracy also slightly decreased as a function of number for the first three numbers (OR = 0.45, CI = [0.38, 0.55], Z = −8.04). In the second segment, enumeration accuracy declined more steeply as a function of number (OR = 0.54, CI = [0.40, 0.73], Z = −3.9). An interaction between disparity condition and number (4-6) in the second segment indicated that enumeration accuracy was initially higher in the second segment during binocular disparity (OR = 1.44, CI = [1.13, 1.86], Z = 3.00). This result is important as it shows that during binocular disparity enumeration remains accurate longer than without disparity. In addition to modelled results, this result is also clearly seen in the observed data: Bonferroni corrected pairwise t tests between the disparity conditions at each number confirmed that a statistically significant difference was only present at number four (difference 8.5 percentage units, p = .002; other comparisons: uncorrected p values ≥ .37). Random-effect variation related to individual differences between participants was SD = 0.80 for intercept, and SD = 0.25 for number.

Reaction times were also best modelled by a two-segment model rather than a corresponding one segment linear model (χ ^2^ = 354.1, df = 37, p < .0001). Reaction times slightly increased for the first three numbers (β = 13.14 ms, CI = [6.46, 19.81], t = 3.86; intercept: β = 616.03 ms, CI = [596.01, 636.06], t = 60.30). For the second segment enumeration times increased more steeply (β = 93.46 ms, CI = [73.299, 105.62], t = 10.85). Thus, whereas binocular disparity increased enumeration accuracy, it did not influence the speed of correct responses. Participant-wise random-effect variation taken into account in the model was as follows: Intercept (SD = 61 ms), number in subitizing range (5 ms), intercept of the second segment (46 ms), number in the second segment (72 ms), presense of disparity (11 ms), number × disparity (first segment: 8 ms; second segment: 86 ms) and second segment intercept × disparity (61 ms).

### Experiment 2: Cube stimuli

Experiments 1 suggests that when viewing natural scenes binocular disparity may increase visual capacity without influencing the speed of visual processing. Experiment 2 was designed to test whether these effects are larger when the stimuli contain very few other depth cues than binocular disparity. In Experiment 2 we employed geometric cubes presented against a distant sky background as the stimuli to be enumerated (see Figure 1). Here other depth cues (than binocular disparity) are minimized whereas the natural scene stimuli employed in Experiment 1 contains various depth cues (e.g. familiar objects in predictable environments, occlusion, shading). We expected this manipulation to enhance the effect binocular disparity has on enumeration performance.

The results are shown in Figure 3. Depth ratings were best fitted by a two-segment model where the first segment included numbers 1-3 (χ ^2^ = 27.3, df = 4, p < .0001). Unlike in Experiment 1, participants’ subjective experience of depth was not modulated by binocular disparity. For numbers 1-3, mean depth rating was 1.90 (β = 1.90, CI = [1.80,2.00], t = 37.6), and for numbers 4-6 depth ratings increased by 0.24 (β = 0.24, CI = [0.18, 0.31], t = 7.3) as a function of number. Participant-wise variation (SD) was 0.32 units in the intercept, and 0.058 in the slope.

**Figure 3:**
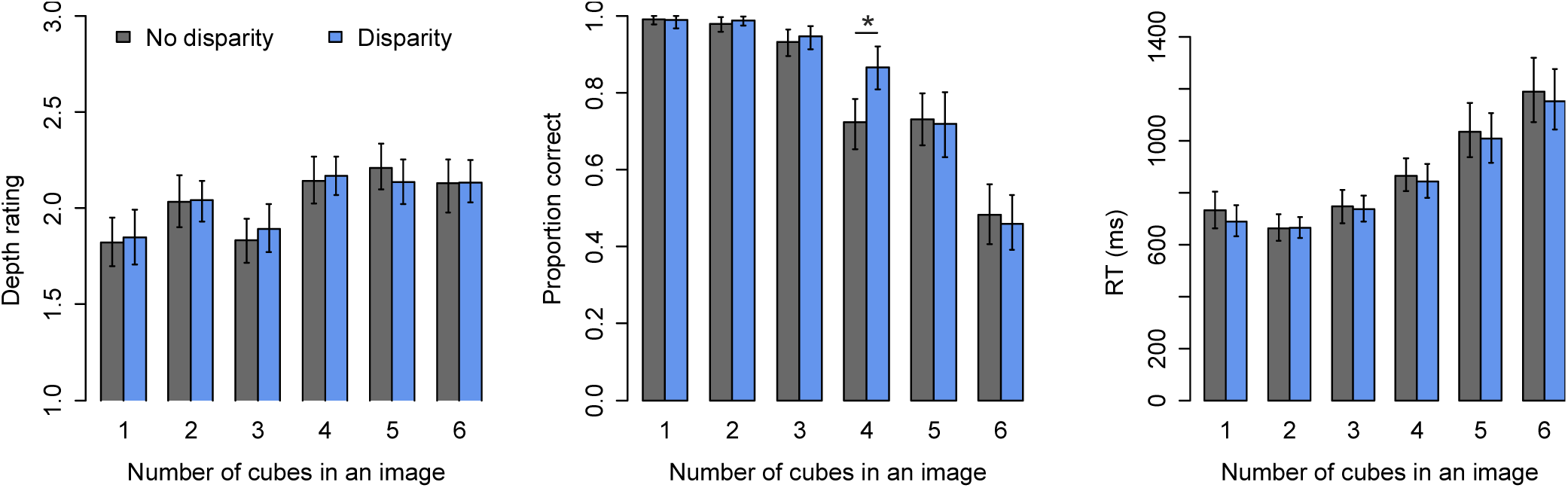
Results of Experiment 2. Error bars show 95% CIs.

The accuracy of enumeration in Experiment 2 was very similar to the results obtained in Experiment 1 with scenes as stimuli. Enumeration accuracy was best fit by a two-segment model (χ ^2^ = 15.1, df = 1, p < .0001). In the first segment, enumeration accuracy decreased a function of number in the first segment (OR = 0.24, CI = [0.18, 0.39], Z = −9.6), and enumeration accuracy was lower when binocular disparity was eliminated (OR = 0.61, CI = [0.43, 0.87], Z = −2.7). Crucially, the second segment also revealed an interaction between number and disparity condition (OR = 1.42, CI = [1.09, 1.84], Z = −2.6): Enumeration remains accurate longer with binocular disparity. Bonferroni corrected t tests at each number comparing the two disparity conditions verified that enumeration accuracy was reduced without binocular disparity (difference 14.3 percentage units; p = .0048) only at number four (other uncorrected p values ≥ .48). All in all, these results with cube stimuli replicate the results obtained with scenes concerning enumeration accuracy. Participant-wise random variations associated with intercept was SD = 1.27, and with number SD = 0.79.

In Experiment 2 reaction times were delayed by a constant of 27 ms when binocular disparity was removed (see Fig 3; β = 27.04 ms, CI = [8.08, 45.99], t = 2.7). That is, with cubes as stimuli, enumeration was faster and more accurate with disparity than without it. Otherwise, the reaction times followed a similar two-segment pattern (χ^2^ = 310.2, p < .0001) as previously. In the first segment reaction times remained constant (β = 693.2 ms, CI = [647.66, 738.81], t = 29.8). During the second segment reaction times increased as a function of number (β = 175.8 ms, CI = [130.71, 220.82], t = 7.6). Participant-wise random-effect variation taken into account in the model was as follows: Intercept (SD = 161 ms), number in subitizing range (47 ms), intercept of the second segment (48 ms), number in the second segment (140 ms), presense of disparity (28 ms), number × disparity (first segment: 18 ms; second segment: 95 ms) and second segment intercept × disparity (70 ms).

## Discussion

Different theories concerning the evolution of frontal eyes have in common the assumption that binocular disparity enables the animals to better discriminate objects from the background 24. The present study examined if binocular disparity aids the enumeration of objects in cluttered, naturalistic scenes in immersive virtual reality. Enumeration paradigm was employed because we were interested if the influence of binocular disparity showed set-size dependent effects. The effects of binocular disparity on objective visual performance (enumeration accuracy and reaction times) and subjectively reported experience of depth were examined. Our results show that even very slight binocular disparity, one that is not noticed by the participant and does necessarily produce a concurrent enhancement of consciously perceived depth, can influence rapid object individuation performance. However, notably our results suggest that the effect of binocular disparity on visual enumeration accuracy and reaction times is very limited: in our study the effect was only observed specifically when four objects were enumerated. Hence, consistent with Valsecchi et al. 12, our results suggest that disparity does not produce a strong general benefit for visual object individuation.

However, the results of the two experiments also reveal a systematic effect at the the “limit” of perceptual capacity. With binocular disparity, the accuracy of enumeration was increased when the scene contained four target individuals, indicating that binocular disparity enhanced the accuracy of perception right at the limit of subitizing capacity. Because disparity only affected enumeration accuracy when the scene contained four items, one may ask if our observation is reliable. Given that virtually identical results were obtained in both experiments using different stimuli, it is unlikely that the observation was caused by chance. Moreover, both experiments had a relatively large sample size and the effect observed was very systematic across individual participants (Bonferroni corrected p values < .005). At number four, disparity increased enumeration accuracy on average by 8.5 and 14.2 percentage units in Experiments 1 and 2, respectively. About similar improvement due to binocular disparity (13%-units on average) was observed for detection of an object in front of a flat textured background 9. The size of the improvement we observed is also roughly similar to the improvement observed when experienced video-game players are compared to non-gamers – as in the present study, the observed increase in enumeration accuracy was about 10 percentage units, and began at number four 47,48. Our findings suggest that in situations where multiple objects compete for perceptual resources, binocular disparity may increase that capacity of perception. This increased performance around the limit of perceptual capacity may have helped animals to better individuate objects (e.g. fruit, prey, predators) that are embedded among other objects (e.g. plants and branches) in complex visual environments 23. Stereoscopic perception constrains the allocation of selective visual attention 49,50, and as subitizing is known to depend on attentional resources 31, the improved enumeration performance under binocular disparity could be due to faster deployment of attention to the objects to be enumerated.

What mechanism could explain that the effect of disparity was observed specifically at single number? Capacity limits in perception have been suggested to result from competition between objects in representational maps: adjacent items inhibit each other and compete for limited “cortical real estate” 51,52. When this competition/inhibition between objects becomes too high, the capacity to individuate objects fails. We suggest that binocular disparity counteracts competition between different objects within the representational maps, enabling the visual system to more efficiently process the objects. More specifically we suggest that when binocular disparity is present, the objects are processed by bisparity-selective neurons, found at multiple visual cortical areas 53,54,37, which reduces competition between assemblies of neurons representing different objects. In other words, when the neurons of the visual system can represent the objects in the scene in three-dimensions (with the aid of visual disparity), the representational maps contains “an extra dimension” so to speak. As the objects are represented by an assembly of neurons which utilize information from binocular disparity, the competition between adjacent objects is reduced (when compared to the situation where disparity is not present). If this type of competition between objects in representational maps underlies perceptual capacity limits 51,52, the advantage provided by disparity would only occur around the limits of perceptual capacity. When only few objects are processed by the representational maps, binocular disparity is not required to individuate the objects. At the other extreme, when the number of objects is clearly higher than the capacity of the system, the limited advantage provided by binocular disparity would not make a difference. However, when the amount of information is just above the limit of perceptual capacity limits, the advantage provided by disparity may reduce competition between objects enough to benefit behavior.

In addition to the advantage observed for enumeration accuracy, disparity also sped-up reaction times by 27 ms on average when geometric cube stimuli were used. While the absolute size of this effect on reaction times is relatively small, it may represent a significant advantage in situations where fast visually-guided behavior is important (e.g. see Julesz, 1971, for similar argument). However, given that this effect was only observed when cube stimuli were used, the present results suggest that this benefit may not generalize to real-world situations. The reader should also note that logically one could have also expected to observe longer reaction times under binocular disparity, as integrating the two different viewpoints of the eyes is a complicated task that can be assumed to require additional processing time. Reaction times were enhanced only in Experiment 2 possibly because other depth cues were minimized with the use of geometric shapes.

Interestingly, our results show that the functional advantage produced by binocular disparity does not require an enhanced conscious perception of depth, suggesting that the mechanisms that produce the functional advantage are somewhat independent from the processes that produce the conscious experience of depth often associated with disparity (Exp 2). Similarly, the results of Experiment 1 show that binocular disparity may produce a clear enhancement in subjectively experienced stereopsis without leading to similar behavioral benefits. The fact that disparity only enhanced the subjective perception of depth in Experiment 1 suggests that subjectively experienced stereopsis is calculated from binocular disparity together with monocular depth cues. The dissociation between enhanced subjective perception of depth and the functional advantage provided by binocular disparity is in concert with recent theoretical proposals that emphasize the difference between functional and subjective forms of stereopsis 3, and with the fact that binocular disparity is not necessary for subjective sensation of depth 35. Moreover, binocular disparities can elicit vergence eye movements in the absence of subjective perception of depth 36. While vergence eye movements and the behavioral effects observed in the present study could be mediated by disparity-selective neurons in V1, these neurons are unlikely to account for subjective perception of depth 37. We suggest that while the functional advantage produced by binocular disparity is likely produced by following rapid computations (possibly in V1; 36,37), the encoding of subjective experience of depth requires longer processing times and integration of information across different cortical areas 55.

The present study also has important limitations. First, further studies are required to replicate the present findings with other stimuli. Future experiments should aim to control and vary stimulus configurations systematically to examine how various stimulus features (density, field-of-view etc.) affect the results, in addition to systematically controlling and varying binocular disparity. In the present study, it is not clear, for example, whether the increase in depth rating as a function of number was caused by mere presence of more objects, or if it had to do with changes in the binocular disparities of those objects. Moreover, while Experiment 1 in the present study included unrelated objects (e.g. trees, rocks), in Experiment 2 the cubes were presented against a distant background. Second, while we aimed to use naturalistic stimuli in Experiment 1, and used virtual reality technology, the stimuli are nevertheless artificial. Use of real stimuli (real individuals in real scenes) could produce different results 27. For instance, in real life, vergence eye movements provide information about distance, whereas with virtual reality headsets the eyes are always focused on the screen surface. Third, future studies should aim to systematically vary object locations, as objects which are closer to the observer (e.g. at reaching distance) are likely to get more benefit from binocular disparity. In the present study the objects were relatively far from the observer. Similarly, there is reason to assume that the size of the advantage produced by binocular disparity would be larger if binocular disparity would be increased by aligning the virtual cameras (i.e. eyes) inward (producing larger disparity), and if the scene would be presented for a longer duration (enabling longer accumulation of visual information).

In conclusion, the present results suggest that binocular disparity may enhance the capacity of visual perception by increasing the number of objects that can be individuated from the background. This effect can arise without concurrent enhancement in consciously experienced depth.

## Acknowledgements

The research was supported by the Finnish Cultural Foundation, and the Academy of Finland (H.R., grant #308533). We thank Iiris Ollikainen for help with data collection.

## Author contributions

H.R. designed the research, analyzed the data, and wrote the article. A.P. programmed the experiments, J.S. and S.K. collected the data.

